## Supplementary information for "Human mitochondrial ferritin exhibits highly unusual iron-O_2_ chemistry distinct from that of cytosolic ferritins"

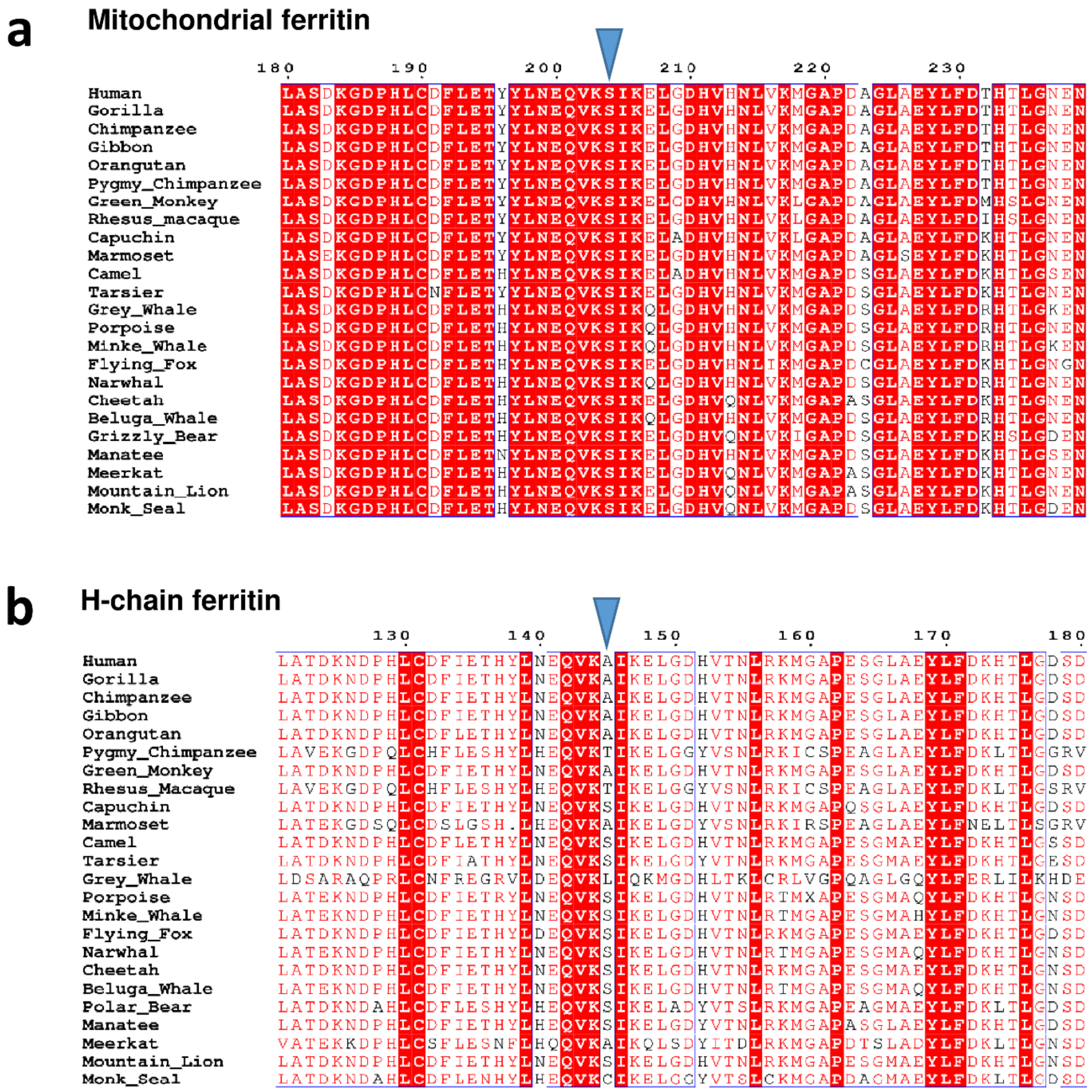


**Figure S1.** Peptide sequences of (**a**) selected predicted FtMt proteins around sequence position 144 and (**b**) the equivalent sequences of selected cytosolic H-chain ferritins. The blue arrow head indicates the position of a serine (Ser144) that is conserved among mitochondrial ferritins but not their cytosolic equivalents. Note that the numbering for the FtMt sequence refers to the full length protein, including the targeting sequence.

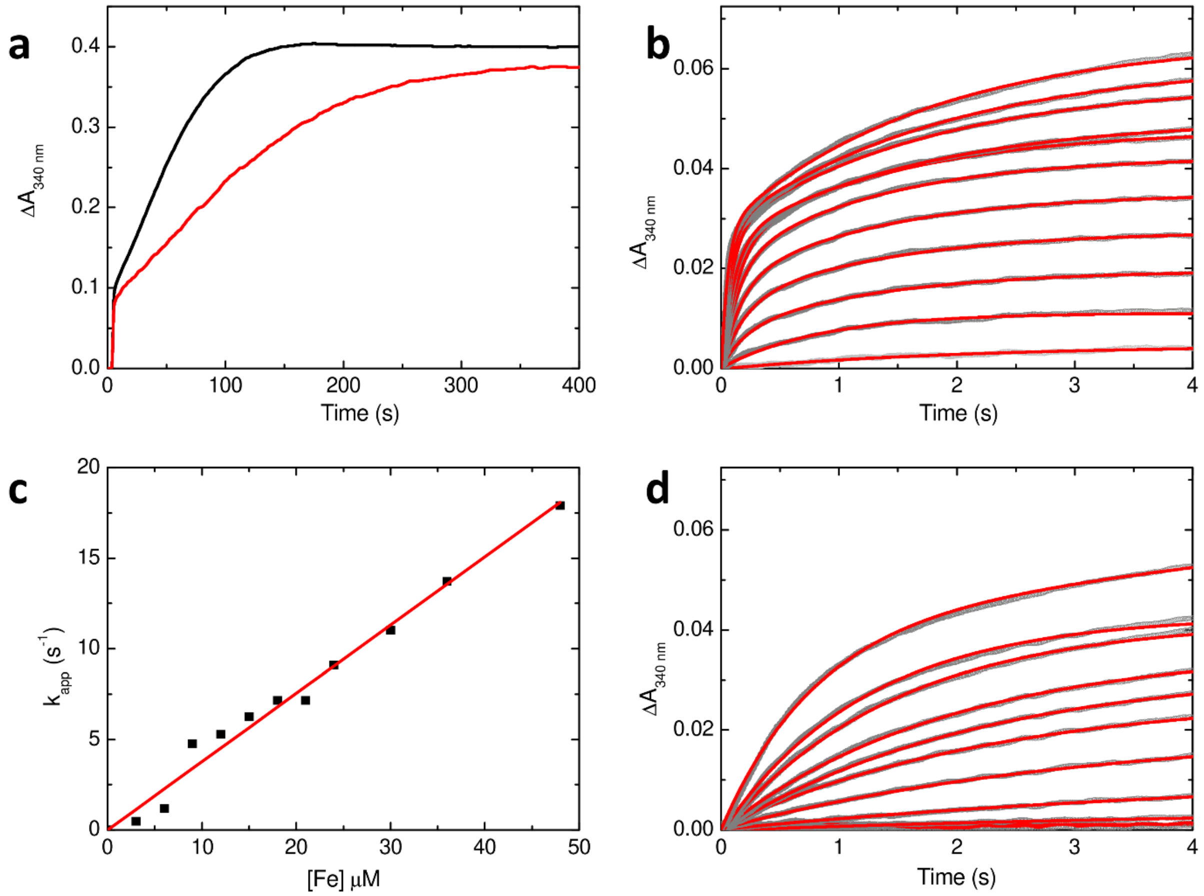


**Figure S2.** Iron oxidation by wild-type and variant Y34F HuHF. (**a**) Increase in absorbance at 340 nm following the aerobic addition of 400 equivalents of $Fe^{2+}$ to 0.5 μM wild-type (black) or variant Y34F (red) HuHF.(**b**) Equivalent data following aerobic mixing of wild-type HuHF and $Fe^{2+}$ leading to final protein concentration of 0.5 μM and iron concentrations between 3 and 48 μM. Data points represented by open grey circles and fits to mono or biexponential functions (as required) by red traces. (**c**) Dependence of the rapid rate of iron oxidation (as apparent first-order rate constants) extracted from the fits in (b) on the concentration of $Fe^{2+}$, yielding a second order rate constant of $3.8 \times 10^5$ $M^{-1}$ $s^{-1}$. (**d**) as (b) but for variant Y34F HuHF.

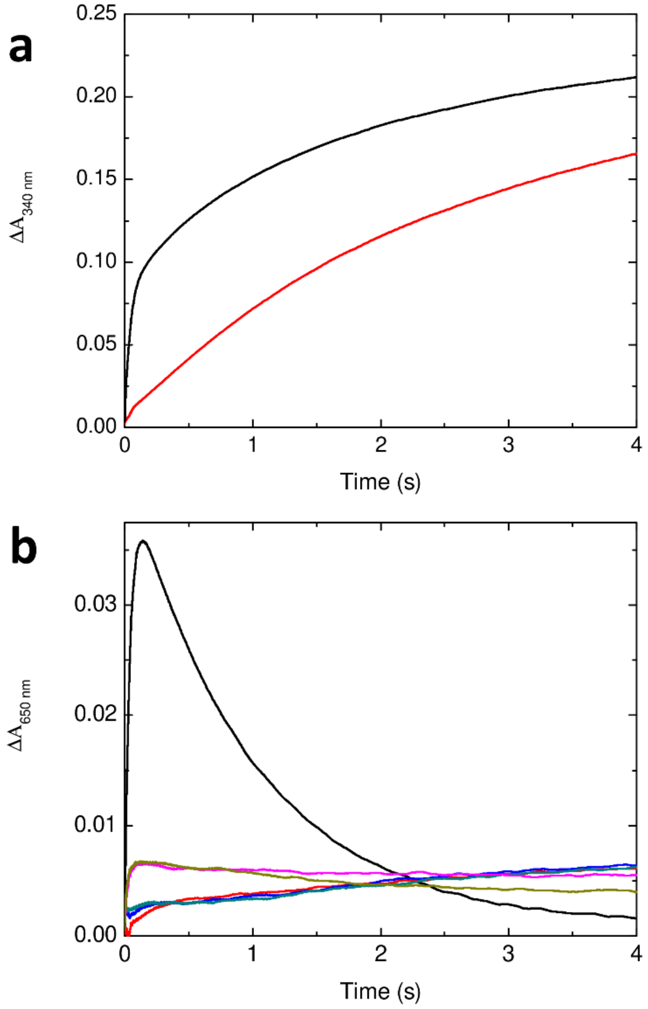


**Figure S3.** The Y34F substitution slows the rate of $Fe^{2+}$ oxidation at FtMt ferroxidase centres. (**a**) Increase in 340 nm absorbance as a function of time following the mixing of 8.33 μM anaerobic wild-type (black) or variant Y34F (red) FtMt containing 48 $Fe^{2+}$ per cage with an equal volume of buffer containing 250 μM dissolved $O_2$ at 10 ˚C. (**b**) Absorbance change at 650 nm as a function of time for samples equivalent to those in (a) and variant Y34F at 15 (blue) 20 (teal) 25 (magenta) and 30 (dark yellow) ˚C.

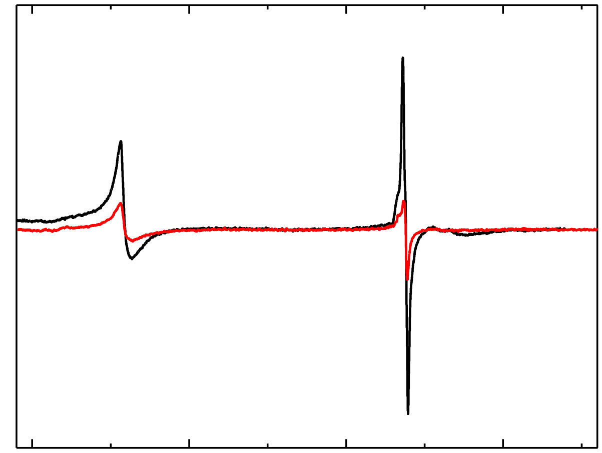


**Figure S4.** Lack of MVFC formation in the reaction cycle of cytosolic human H-chain ferritin. The EPR spectrum of human H-chain ferritin following anaerobic incubation with 48 equivalents of $Fe^{2+}$ prior to mixing with oxygenated buffer giving final concentrations of 4.17 µM human H ferritin, 200 µM $Fe^{2+}$ and 50 µM $O_2$ (black trace) and 4.17 µM variant Y34F human H-chain ferritin aerobically mixed with 300 µM $Fe^{2+}$ (red trace). Samples were frozen approximately 10 s after initiating the oxidation reactions. The spectra lack the signals at magnetic fields greater than 3400 Gauss characteristic of MVFC formation, which are evident in equivalently treated samples of mitochondrial ferritin (see Fig. 4 of the main paper).

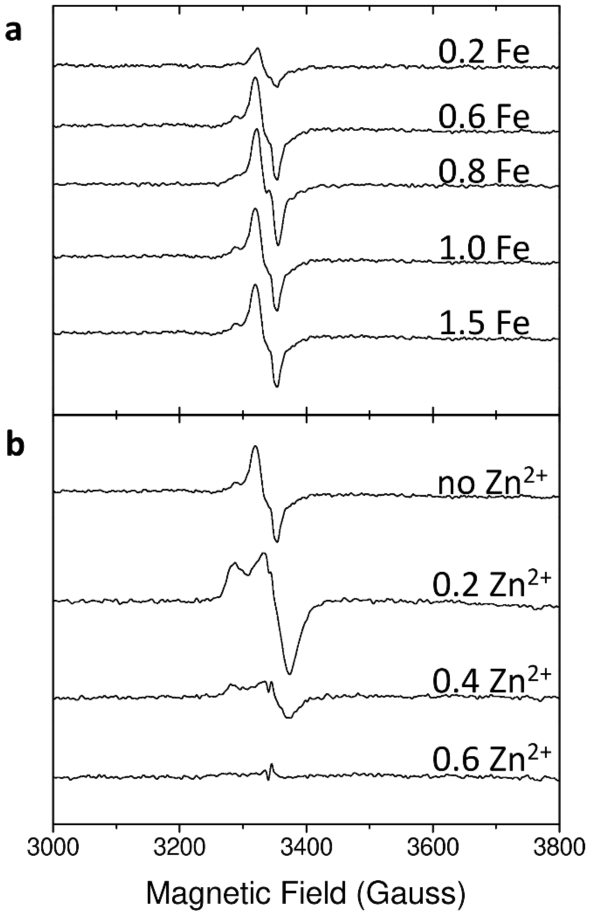


**Figure S5.** The effect of $Fe^{2+}$:protein stoichiometry and pre-incubation with $Zn^{2+}$ on $Fe^{2+}$ oxidation catalyzed by FtMt. The dependence of the rate of $Fe^{2+}$ oxidation at the FC of variant Y34F on $Fe^{2+}$ concentration suggested that $Fe^{2+}$ binding to the catalytic site is impaired in this protein (Fig. 2d). This raised the possibility that observation of an MVFC during the $Fe^{2+}$ oxidation was simply due to some FC sites being partially occupied by $Fe^{2+}$, with subsequent transfer of an electron to a di-$Fe^{3+}$ FC. Alternatively, unreacted $Fe^{2+}$ remaining in solution may be able to displace one $Fe^{3+}$ from FCs (demonstrated to lead to MVFC formation in HuHF [1]. The consequence of partial occupancy of wild-type FtMt FCs was probed by EPR following the aerobic exposure ($O_2$ in excess) of wild-type FtMt to $Fe^{2+}$ at concentrations below that required to fill all FC sites. (**a**) shows EPR spectra of wild-type FtMt frozen 10 s after aerobic mixing with the indicated stoichiometry of $Fe^{2+}$ per FOC iron-binding site. Additionally, experiments were performed in which 3 equivalents of $Fe^{2+}$ per FC were added following pre-incubation with variable amounts of $Zn^{2+}$, a potent inhibitor of ferritin activity that acts by binding at FC sites preventing the binding of $Fe^{2+}$ substrate [2]. (**b**) shows EPR spectra of wild-type FtMt pre-incubated with the indicated stoichiometry of $Zn^{2+}$ per FOC iron-binding site prior to aerobic mixing with 1.5 $Fe^{2+}$ per iron-binding site and freezing 10 s after initiating the reaction. Data indicate that catalysis of $O_2$-driven $Fe^{2+}$ oxidation by FtMt is suppressed at a stoichiometry of 1.2 $Zn^{2+}$ per FOC (0.6 per FOC iron-binding site). Experiments in both (a) and (b) could result in FCs containing only one $Fe^{2+}$ ion, with potential for electron transfer to a di-$Fe^{3+}$ FC. In (b), pre-incubation with $Zn^{2+}$ also results in unreacted $Fe^{2+}$ remaining in solution and able to displace $Fe^{3+}$ from the di-$Fe^{3+}$ sites expected to form in the initial 500 ms of the reaction. However, the resulting spectra contained no evidence of MVFC formation.

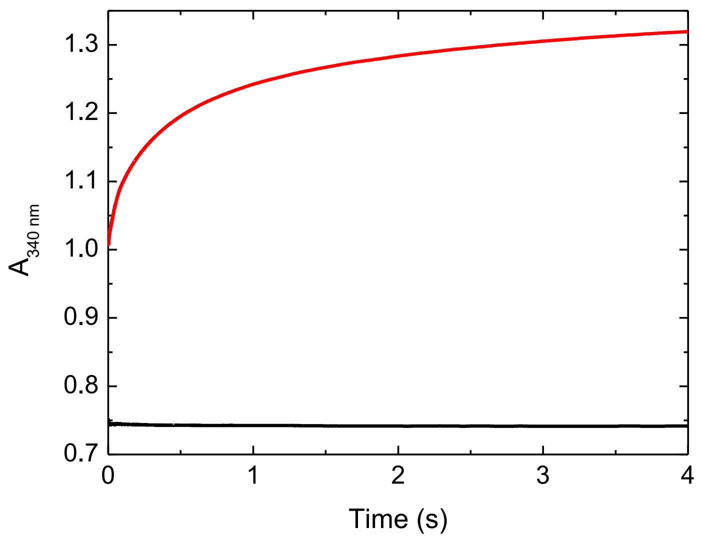


**Figure S6.** Reaction of FtMt in the MVFC form with $O_2$. Mixing of 60 µL of a 50 mM $Fe^{2+}$ solution with 3 mL of 20.8 µM wild-type FtMt within a gas-tight Hamilton syringe led to a solution with absorbance at 340 nm of 0.7. Rapid stopped-flow mixing with an equal volume of anaerobic MES pH 6.5 resulted in little change in absorbance (black trace), whilst mixing with aerobic buffer (red trace) resulted in an increase in absorbance to approximately 1.0 within the dead time of the instrument followed by a slower increase to a final value of approximately 1.3.

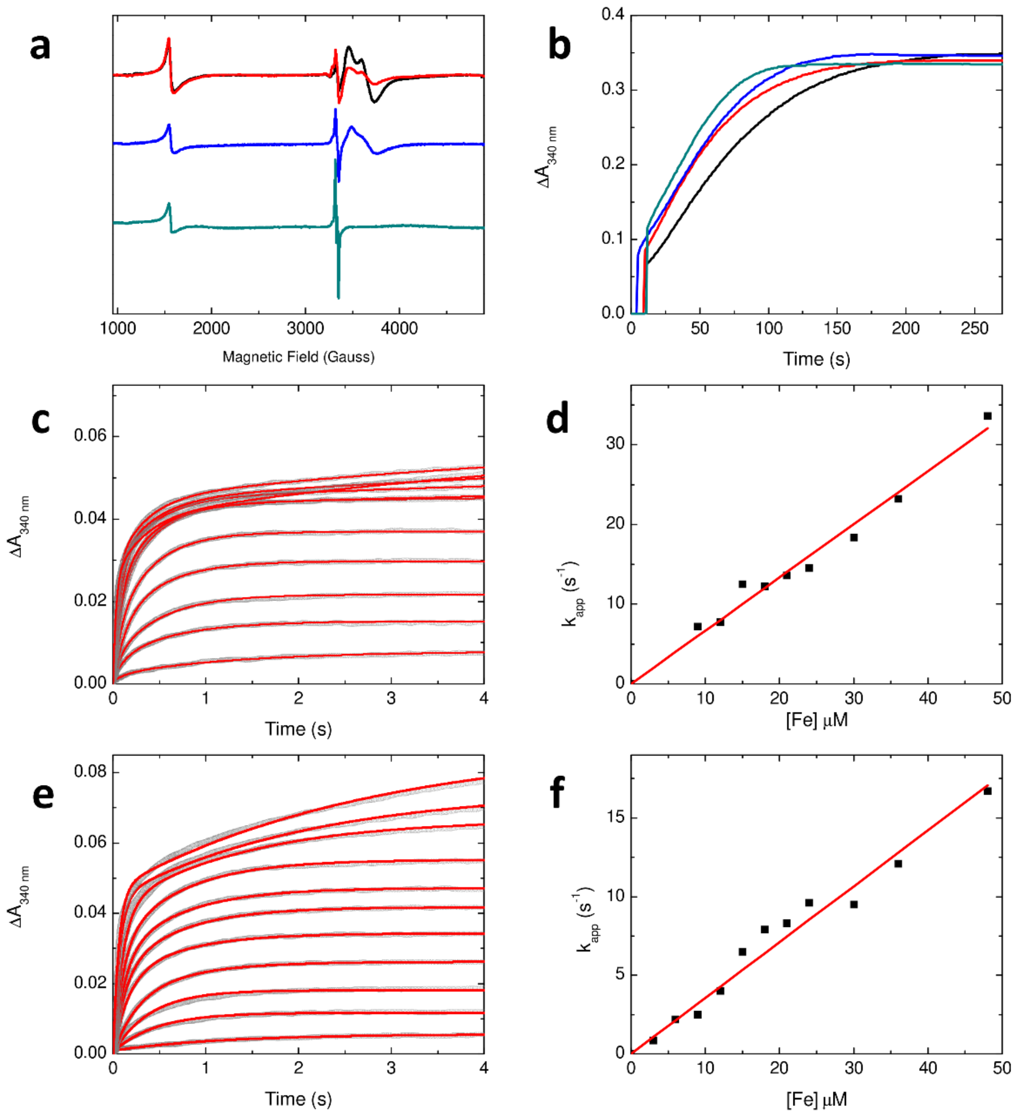


**Figure S7.** (**a**) EPR spectra of the products of FtMt activity under limiting $O_2$. The EPR spectrum of 300 μL of a 20.8 μM solution of wild-type FtMt frozen 10 s after aerobic mixing with 6 μL of a 50 mM $Fe^{2+}$ solution (50 $Fe^{2+}$ per FtMt, $O_2$ sufficient to oxidize 50% of added $Fe^{2+}$, black trace). Red trace shows the equivalent spectrum following the equal volume mixing of 41.6 μM protein and 2 mM $Fe^{2+}$ solutions (50 $Fe^{2+}$ per FtMt, $O_2$ sufficient to oxidize 50% of added $Fe^{2+}$). We note that the yield of MVFC for the latter was reduced compared to the addition of a small volume of 50 mM $Fe^{2+}$. This reduction may have been due to the introduction of further $O_2$ to the solutions during mixing. Blue and cyan traces are equivalent to the black trace but using FtMt S144A and HuHF A144S in place of wild-type FtMt. (**b**) mineralization activity of FtMt S144A (red trace) with wild-type protein in black for comparison, and of HuHF variant A144S (blue trace) with wild-type protein in cyan for comparison. (**c**) Increase in 340 nm absorbance following stopped-flow mixing of FtMt S144A with $Fe^{2+}$ solutions of varying concentration to give final concentration of 0.5 μM protein and 3-48 μM $Fe^{2+}$. Data shown as open grey circles and fits to mono or bi exponential functions (as required) as red traces. (**d**) Dependence of the rapid rate of iron oxidation extracted from the fits in (c) on iron concentration. (**e**) and (**f**) as for (c) and (d) but using HuHF A144S.

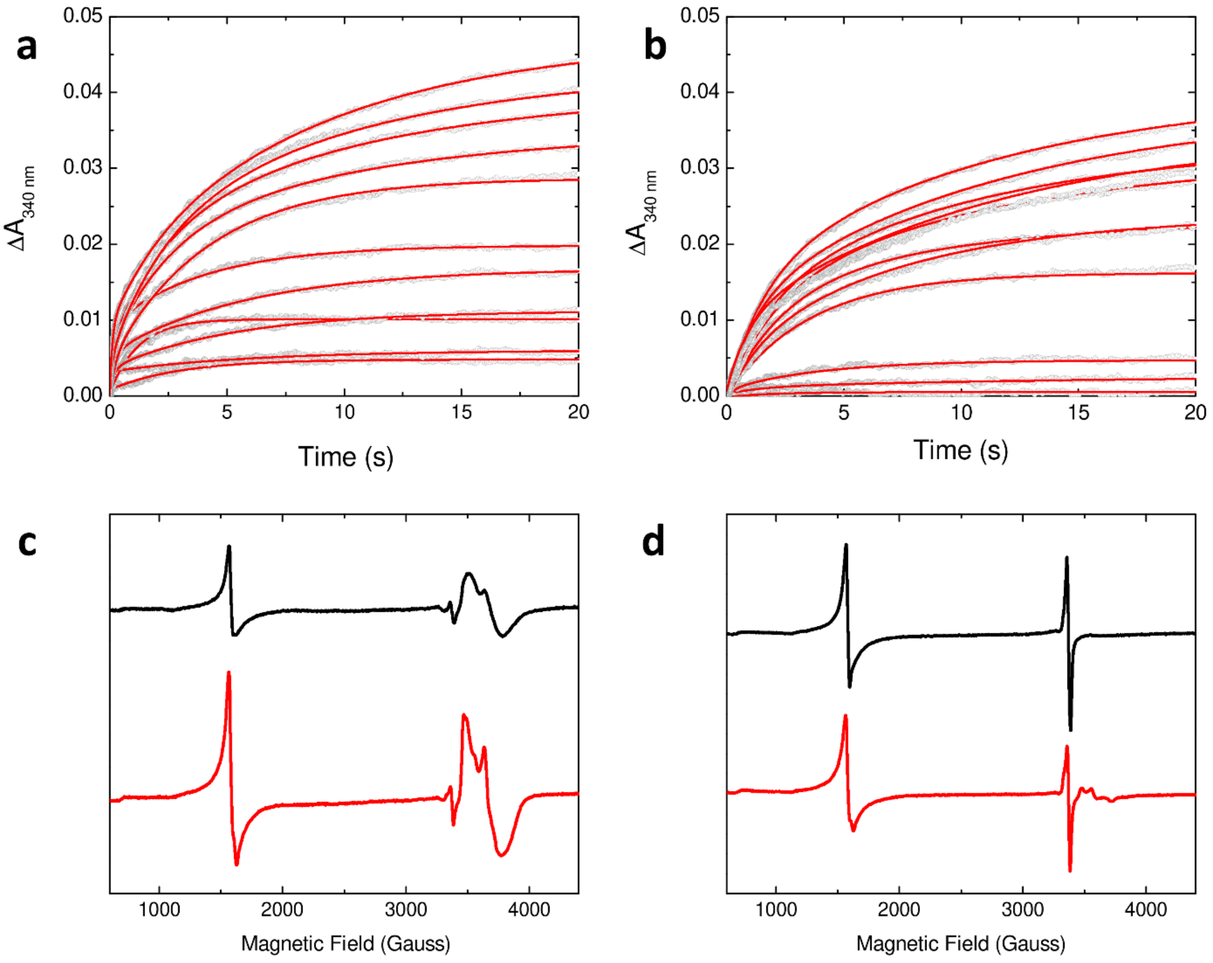


**Figure S8.** Reactivity of FtMt towards $H_2O_2$. The increase in 340 nm absorbance as a function of time following the anaerobic equal volume mixing of 1 μM solutions of (**a**) wild-type and (**b**) Y34F FtMt containing 500 μM $H_2O_2$ with $Fe^{2+}$ solutions of concentration 6-96 μM. Open gray circles indicate data points, red traces represent fits to mono- or bi-exponential fits as required. (**c**) EPR spectra of wild-type (black) and Y34F (red) FtMt anaerobically incubated with 300 μM $Fe^{2+}$ prior to mixing with 100 μM $H_2O_2$ and frozen approximately 10 s after $H_2O_2$ addition. (**d**) as (a) but using 500 μM $H_2O_2$. The data confirm that peroxide driven oxidation of iron by FtMt is much slower than the reaction with oxygen but in both cases variant Y34F forms significant quantities of MVFC whilst the wild type protein only does so when iron is present in excess of oxidant, regardless of co-substrate.

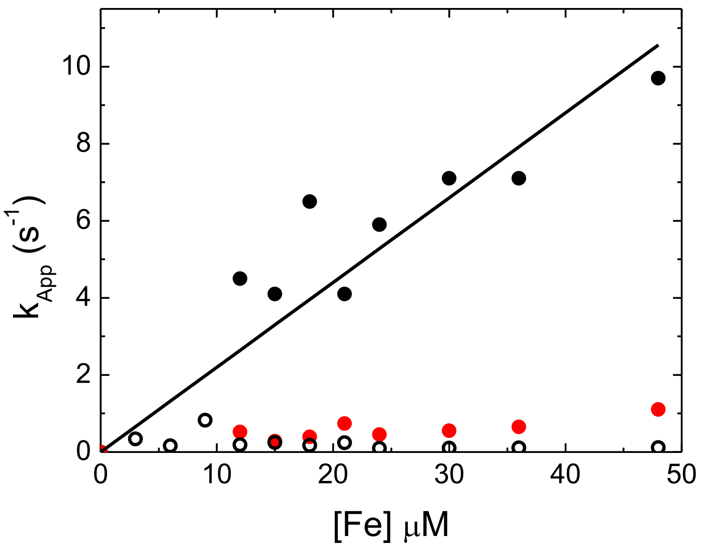


**Figure S9.** Reaction of FtMt with $H_2O_2$ is complex. Fitting of kinetic data shown in Fig. 5 of the main paper required a bi-exponential function for wild-type FtMt. The amplitude of the initial, fast phase was small, accounting for only ~10% of the total amplitude. A plot of the first order rate constant associated with this fast phase is shown (black circles). The small amplitude of this phase resulted in a greater uncertainty in the apparent first order rate constants, but fitting of their dependence to a linear function yielded a second order rate constant of 2.2 x$10^5$ $M^{-1}$ $s^{-1}$. This is similar to that determined from a similar analysis of the 10-fold higher amplitude of the rapid phase observed upon reaction with $O_2$, which also was $Fe^{2+}$-dependent. This indicated that the rate-limiting step of the reaction was $Fe^{2+}$ binding. Although it is unclear what the small increase in amplitude observed upon mixing of $Fe^{2+}$ with FtMt in the presence of $H_2O_2$ is due to, it was also rate-limited by $Fe^{2+}$ binding. Rate constants associated with the second, slower phase are also plotted (black open circles), consistent with a much slower reaction that is independent of $Fe^{2+}$ concentration. A similar analysis of data acquired for Y34F FtMt revealed that there was no fast phase and so a single exponential function was used for fitting. The resulting rate constants are plotted above (red circles), indicating that this slow phase is independent of $Fe^{2+}$ concentration. These data suggest that the conserved Tyr may also be important in activating the FC of FtMt to react with $H_2O_2$, but the effect of the absence of the Tyr is much less severe than for the reaction with $O_2$.

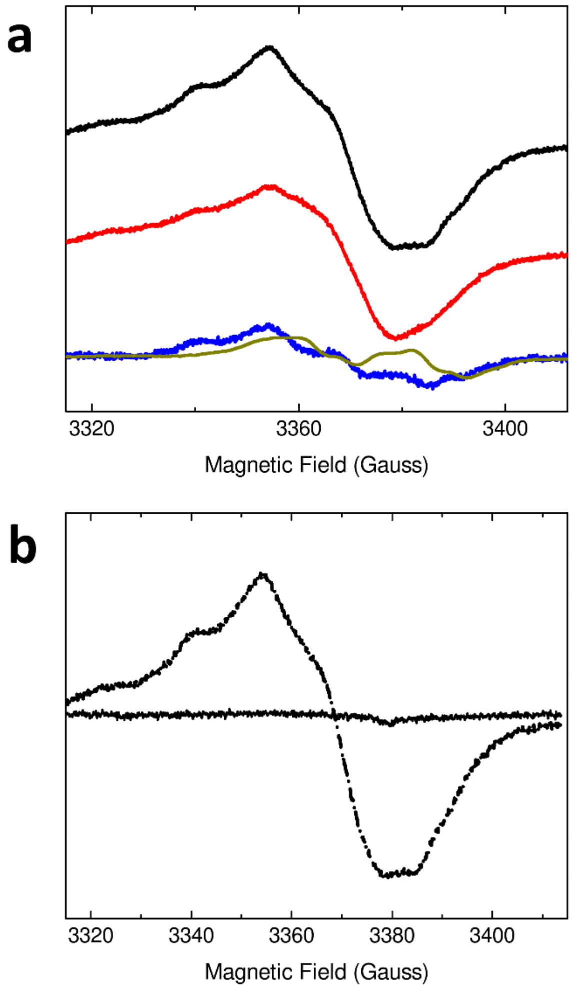


**Figure S10.** Radical formation during the oxidation of $Fe^{2+}$ by $H_2O_2$. (**a**) Spectra of wild-type (black) and Y34F (red) FtMt anaerobically incubated with 300 μM $Fe^{2+}$ prior to mixing with 500 μM $H_2O_2$ and frozen approximately 10 s after $H_2O_2$ addition. Blue trace shows the difference spectrum between the protein-based radical formed on the wild-type and Y34F proteins; dark yellow trace is the equivalent plot for the $O_2$-driven reaction. (**b**)The EPR spectrum of wild-type FtMt inhibited by pre-incubation with 60 equivalents of $Zn^{2+}$ prior to anaerobic incubation with 300 μM of $Fe^{2+}$ and then 500 μM $H_2O_2$ (solid black trace). Dashed black trace shows the equivalent spectrum recorded in the absence of $Zn^{2+}$ demonstrating that formation of the protein-based radical is dependent on FOC activity. Samples were frozen approximately 10 s after the addition of $H_2O_2$.

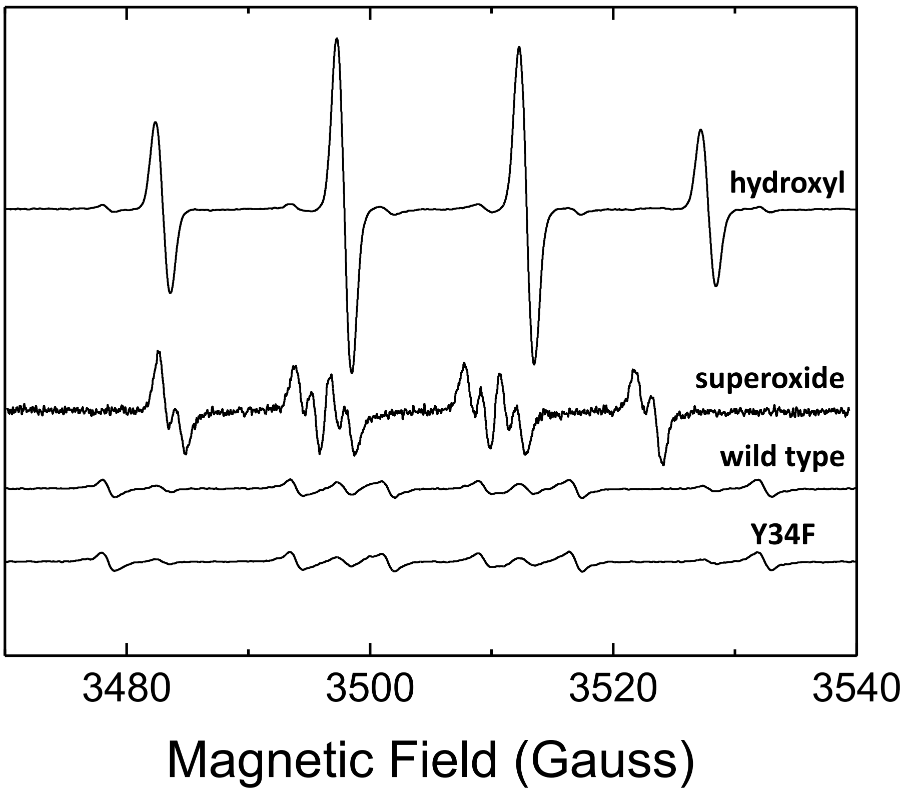


**Figure S11.** $O_2$ reduction during $Fe^{2+}$ oxidation catalysed by FtMt. Amplex Red assays were conducted with an $Fe^{2+}$ concentration of 9.6 μM, giving a theoretical maximum yield of 4.8 μM $H_2O_2$. Protein concentration was varied between 0.2 and 1.0 μM to investigate the effect of partial loading of FCs on the product of $O_2$ reduction. For both wild-type and variant Y34F FtMt, the concentration of $H_2O_2$ released, as judged by reference to a calibration plot constructed using serial dilution of a standard $H_2O_2$ solution, was approximately 1.1 μM, irrespective of the ratio of $Fe^{2+}$ to protein employed. A control experiment in which 1.0 μM FtMt with di-$Fe^{3+}$ FCs was mixed with 5 μM $H_2O_2$ and subjected to the Amplex Red assay also led to detection of 1.1 μM $H_2O_2$, suggesting that the presence of FtMt interferes with $H_2O_2$ detection either by direct reaction or reaction with Complex I of horse radish peroxidase, preventing oxidation of the Amplex Red dye. Radical trapping experiments during $Fe^{2+}$ oxidation catalysed by FtMt were also performed. Shown here are EPR spectra of the hydroxyl and superoxide radical adducts of the spin trap DMPO, generated by incubating the reagent with $Fe^{2+}$ and $H_2O_2$ or xanthine/xanthine oxidase respectively, together with corresponding DMPO spectra following aerobic incubation with either wild-type or variant Y34F FtMt and $Fe^{2+}$. The spectra following incubation with either protein are indistinguishable and demonstrate negligible formation of superoxide in either case. Thus, the data are consistent with the conclusion that peroxide is the major product of $O_2$ reduction for both wild-type and variant proteins.

**Table S1** Iron oxidation by the ferroxidase centres of FtMt[a].

| $Fe^{2+}$/FtMt | Wild-type | | | | Y34F | | | |
|---|---|---|---|---|---|---|---|---|
| | $k_{rapid}$ ($s^{-1}$) | $A_{rapid}$ | $k_{slow}$ ($s^{-1}$) | $A_{slow}$ | $k_{rapid}$ ($s^{-1}$) | $A_{rapid}$ | $k_{slow}$ ($s^{-1}$) | $A_{slow}$ |
| 6 | 0.89 | 0.0030 | - | - | - | - | 0.19 | 0.0006 |
| 12 | 1.90 | 0.0103 | - | - | - | - | 0.15 | 0.0040 |
| 18 | 2.65 | 0.0188 | - | - | 2.01 | 0.0025 | 0.19 | 0.0074 |
| 24 | 2.71 | 0.0244 | - | - | 1.65 | 0.0033 | 0.19 | 0.0091 |
| 30 | 3.08 | 0.0305 | - | - | 1.80 | 0.0052 | 0.17 | 0.0109 |
| 36[b] | 3.22 | 0.0346 | - | - | 1.69 | 0.0072 | 0.14 | 0.0129 |
| 42[b] | 3.68 | 0.0383 | - | - | 1.76 | 0.0093 | 0.11 | 0.0163 |
| 48[b] | 3.94 | 0.0395 | - | - | 1.87 | 0.0095 | 0.11 | 0.0183 |
| 60 | 5.15 | 0.0397 | 0.07 | 0.0088 | 2.36 | 0.0114 | 0.09 | 0.0238 |
| 72 | 6.25 | 0.0390 | 0.07 | 0.0128 | 3.38 | 0.0112 | 0.10 | 0.0269 |
| 96 | 8.33 | 0.380 | 0.11 | 0.019 | 4.99 | 0.0120 | 0.09 | 0.0348 |

[a]Effective first order rate constants and associated amplitudes of the rapid and slow phases of FC catalysed $Fe^{2+}$ oxidation for wild type and variant Y34F FtMt.

[b]At $Fe^{2+}$ loadings of 36 to 48 $Fe^{2+}$/FtMt the wild-type protein exhibited a linear increase in absorbance at 340 nm following the rapid phase of iron oxidation. The amplitude of this increase was extremely small and did not support the extraction of an effective first order rate constant.
